# Exploring the pH-dependent structure-dynamics-function relationship of human renin

**DOI:** 10.1101/2020.10.15.340935

**Authors:** Shuhua Ma, Jack A. Henderson, Jana Shen

## Abstract

Renin is a pepsin-like aspartyl protease and an important drug target for the treatment of hypertension; despite three decades’ research, its pH-dependent structure-function relationship remains poorly understood. Here we employed the continuous constant pH molecular dynamics (CpHMD) simulations to decipher the acid/base roles of renin’s catalytic dyad and the conformational dynamics of the flap, which is a common structural feature among aspartyl proteases. The calculated p*K*_a_’s suggest that the catalytic Asp38 and Asp226 serve as the general base and acid, respectively, in agreement with experiment and supporting the hypothesis that renin’s neutral optimum pH is due to the substrate-induced p*K*_a_ shifts of the aspartic dyad. The CpHMD data confirmed our previous hypothesis that hydrogen bond formation is the major determinant of the dyad p*K*_a_ order. Additionally, our simulations showed that renin’s flap remains open regardless of pH, although a Tyr-inhibited state is occasionally formed above pH 5. These findings are discussed in comparison to the related aspartyl proteases, including *β*-secretases 1 and 2, capthepsin D, and plasmepsin II. Our work represents a first step towards a systematic understanding of the pH-dependent structure-dynamics-function relationships of pepsin-like aspartyl proteases that play important roles in biology and human disease states.

## INTRODUCTION

Renin-angiotensin-aldosterone system (RAAS) is a critical regulator for blood pressure and systemic vascular resistance.^1,2^ As an aspartic protease and a part of the RAAS system, renin cleaves the protein angiotensinogen to generate an inactive decapeptide, angiotensin I, which is further cleaved by the angiotensin converting enzyme (ACE) to produce shorter peptides, including the octapeptide angiotensin II which binds and activates the angiotensin II type 1 (AT_1_) and type 2 (AT_2_) receptors. The primary effects of AT_1_ receptor activation include vasoconstriction and stimulation of aldosterone synthesis and release with subsequent sodium and fluid retention, which tend to elevate blood pressure.^1^ Because of its essential role in the function of RAAS and high specificity for its only known substrate angiotensinogen, renin is an attractive drug target for the treatment of hypertension. Despite the considerable efforts,^3–8^ so far Aliskiren is the only renin inhibitor that has reached the market for the treatment of high blood pressure. Interestingly, ACE2, which is a homolog of ACE, counters RAAS activation by processing angiotensin II,^9^ it also functions as a receptor for SARS coronaviruses.^10^ The potential effect of RAAS inhibitors used among COVID-19 patients who have hypertension has been a topic of intense debate.^10^

Renin belongs to the A1 family of aspartic peptidases, which are also known as pepsin-like enzymes due to the structural similarity to pepsin.^11,12^ This subfamily also includes other monomeric aspartic proteases such as cathepsin D (CatD), *β*-secretases 1 and 2 (BACE1 and BACE2), and plasmepsin II (PlmII). Renin shares a sequence identity level of 46% with CatD, 33% with PlmII, 25% with BACE2, and 24% with BACE1, and the respective sequence similarity levels are 66%, 53%, 47%, and 43% (Figure 1). Similar to these other aspartic proteases, the active form of renin is a 340-residue monomer, which folds to a predominantly *β*-sheet structure, whereby a *β*-hairpin loop known as the flap (Thr80 to Gly90) is located over the active site comprised of an aspartic dyad (Asp38 and Asp226)^13^ (Figure 2).

**Figure 1.**
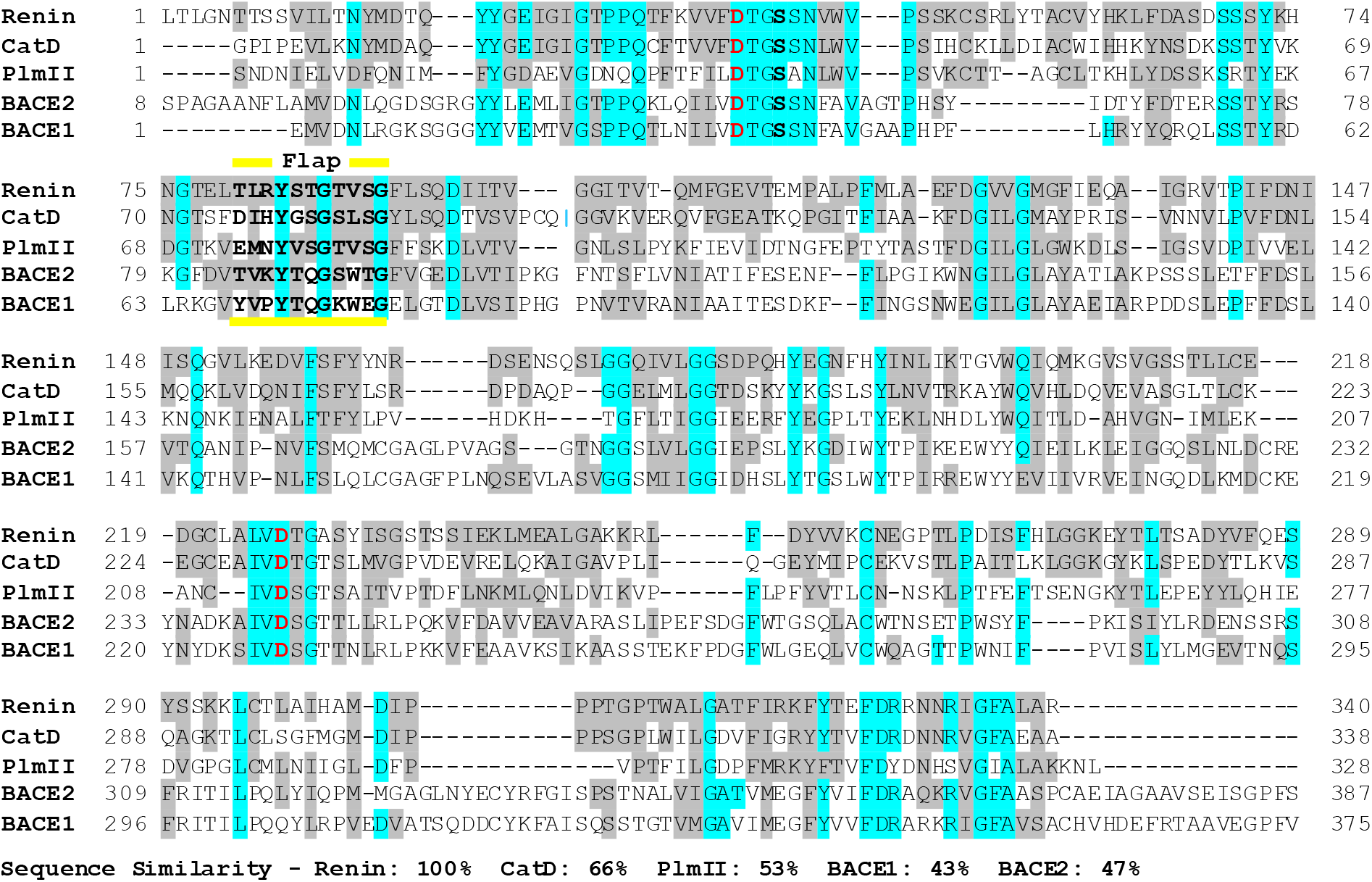
Sequence alignment of renin, cathepsin D, plasmepsin II, BACE1 and BACE2. Conserved and similar residues are highlighted in cyan and gray, respectively. The catalytic aspartic residues are colored red and the sequence of the flap region is indicated in bold font and labeled.

**Figure 2.**
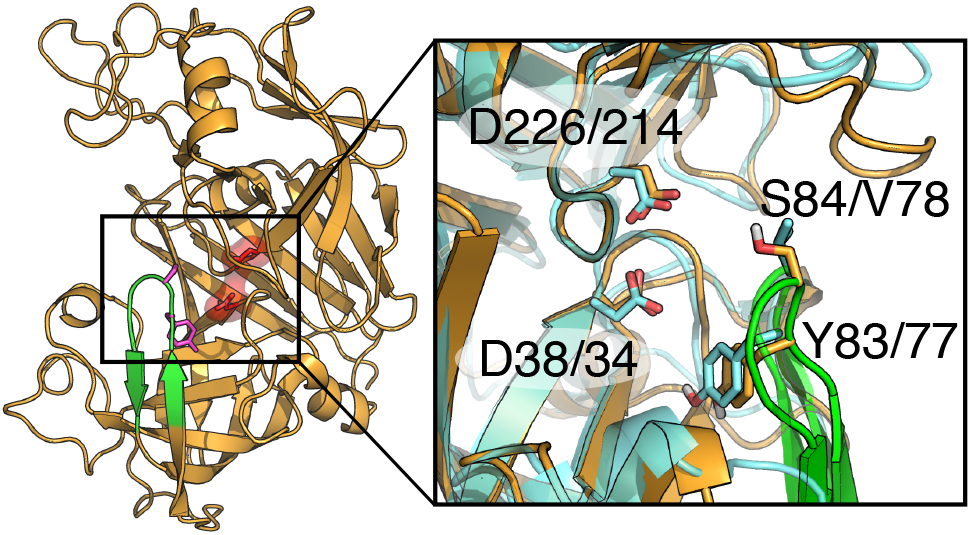
The overall structure of human renin and its substrate binding site. Left: The X-ray crystal structure of human renin (PDB: 2ren^13^). The flap (residues Thr80 to Gly90) and dyad (Asp38 and Asp226) are colored green and red, respectively. Right: A zoomed-in view of the substrate binding site and flap region of renin (orange) overlaid on the X-ray structure of plasmepsin II (cyan, PDB: 1sme^32^). Residues discussed in the main text are labeled.

The catalytic activity of renin is pH dependent; however, unlike a typical pepsin-like aspartyl protease which has a bell-shaped pH profile with an acidic optimum pH, renin’s catalysis occurs in the pH range 5.5–8, with a peak near neutral pH.^14–18^ Interestingly, several experimental studies found that the peak of renin’s pH-activity profile shifts dependent on the substrate.^14–20^ Based on the pH-activity measurements of renin with wild-type and mutant angiotensinogen, it was hypothesized that the active-site residues interact with the His residues at the P2 and P3’ positions, resulting in the unusual optimum pH of renin.^16,17,19,21^ However, the detailed pH-dependent mechanism remains unclear.

The catalytic function of an aspartyl protease is carried out by the catalytic aspartic acids, whereby the general base or nucleophile is deprotonated (lower p*K*_a_) while the general acid or proton donor is protonated (higher p*K*_a_) in the enzyme active pH range. Thus, a mechanistic understanding of the catalytic function starts with the knowledge of the dyad protonation states.^22^ However, existing theoretical and computational studies of renin either do not distinguish the dyad residues or assign protonation states to them in an ad hoc fashion.^23–25^ For example, Calixto et al.^25^ employed a quantum mechanical/molecular mechanical (QM/MM) method to study the catalytic mechanism of renin, but the protonation states of the dyad were assigned (protonated Asp38 and unprotonated Asp226) based on a proposed mechanism of HIV-1 protease. Note, renin and HIV-1 protease belong to different subfamilies of aspartyl proteases, and there is no experimental or theoretical evidence to support the same p*K*_a_ order for the aspartic dyad.

In this work, we employed the hybrid-solvent continuous constant pH molecular dynamics (CpHMD) simulations with pH replica-exchange^26,27^ to determine the catalytic dyad protonation states of renin and characterize the possibly pH-dependent conformational dynamics of the flap. The replica-exchange hybrid-solvent CpHMD method has been previously applied to study several pepsin-like aspartyl proteases BACE I,^28,29^ BACE II,^30^ and CatD^29^ and obtained dyad p*K*_a_ order in agreement with experiments. These studies and another work that examined also other enzymes,^22^ demonstrated that the general base or nucleophile forms more hydrogen bond than the general acid or proton donor and some of the hydrogen bonds are absent in the crystal structure and emerge during proton-coupled conformational sampling. The present CpHMD simulations of renin confirmed this hypothesis. Interestingly, we found that the aspartic dyad residues in renin have opposite catalytic roles as the analogous residues in BACE1, BACE2, and CatD, despite the sequence and structural similarities. To corroborate the finding, we also performed a simulation for PlmII, which belongs to the same pepsin-like protease family. We found that the aspartic dyad residues in PlmII have the same acid/base roles as the analogous ones in renin. The simulation results allowed us to test the experimental hypothesis and understand why renin’s catalytic function occurs in a neutral pH despite the low p*K*_a_’s of the dyad residues. Finally, we found that renin’s flap is open regardless of pH; however, a Tyr-inhibited state due to the formation of a hydrogen bond between Tyr and Asp analogous to BACE1 is formed above pH 5. These findings enable us to begin a journey to systematically understand the pH-dependent structure-activity relationships of pepsin-like aspartyl proteases which play important roles in biology and human disease states.

## RESULTS and DISCUSSION

We performed the pH replica-exchange hybrid-solvent CpHMD simulations starting from the X-ray crystal structure of the apo human renin (PDB: 2ren^13^). 24 independent pH replicas in the pH range 1-9 underwent constant NPT simulations, with an aggregation sampling time of 672 ns. To substantiate the statistical significance of our results, another set of replica-exchange CpHMD simulations was performed starting from the X-ray structure of renin in complex with an inhibitor (PDB: 3sfc^31^) but with the inhibitor removed. The aggregation sampling time was 648 ns. The p*K*_a_’s of the dyad residues in both sets of simulations were well converged (Figure S1). The data from the first 4 ns (per replica) was discarded in the analysis.

### Asp38 is the general base and Asp226 is the general acid

We first examine the pH titration of the aspartic dyad Asp38 and Asp226 in renin. By fitting the residue-specific unprotonated fractions as a function of pH to the generalized Henderson-Hasselbalch equation, we obtained the microscopic p*K*_a_’s of 3.7 and 5.2 for Asp38 and Asp226, respectively (Figure 3a, Table 1). Like many other catalytic aspartyl dyads, ^22^ the carboxylate sidechains of Asp38 and Asp226 are hydrogen bonded to each other in the crystal structure (a minimum oxygen-oxygen distance of 2.9 Å in PDB: 2ren^13^). Therefore, to characterize the coupled titration, we calculated the macroscopic stepwise p*K*_a_’s by fitting the total number of protons of the dyad as a function of pH to a coupled two-proton model (Eq. 2), which resulted in the p*K*_a_’s of 3.2 and 5.3 (Table 1, Figure S2). Compared to the microscopic p*K*_a_’s, the splitting between the stepwise p*K*_a_’s is increased by 0.6 pH units, indicating a small degree of coupling between the titration of Asp38 and Asp226.

**Figure 3.**
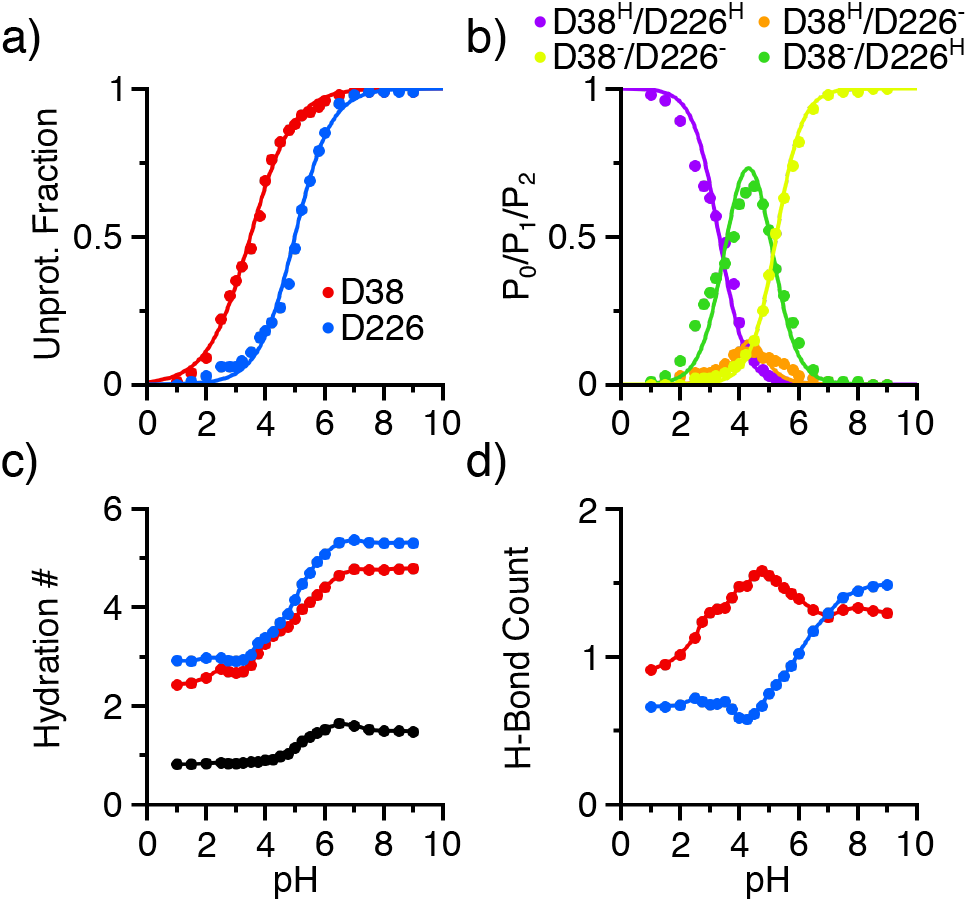
pH titration and pH-dependent properties of the catalytic dyad residues in renin. **a)** Unprotonated fractions of Asp38 (red) and Asp226 (blue) as a function of pH. The lines are the best fits to the generalized Henderson-Hasselbalch equation. **b)** Probabilities of the doubly deprotonated (*P*_0_, yellow), singly protonated (*P*_1_, orange and green), and doubly protonated (*P*_2_, purple) states. **c)** Hydration number of Asp38 (red) and Asp 226 (blue) as well as bridge water (black) between the catalytic dyad residues as a function of pH. **d)** Total number of hydrogen bonds formed by Asp38 (red) and Asp226 (blue) as a function of pH. Data from the simulation run 1 was used.

In order to understand the sequence of titration events, we calculated the pH-dependent probabilities of the four possible protonation states of the dyad. As pH decreases from 7 to 4, the probability of the doubly deprotonated state D38^−^/D226^−^ decreases to nearly zero, while that of the singly protonated state D38^−^/D226^H^ increases to a maximum of about 75% and that of the alternative singly protonated state D38^H^/D226^−^ increases to about 10% (Figure 3b). As pH further decreases to 2, the probabilities of both singly protonated states decrease to nearly zero, while that of the doubly protonated state D38^H^/D226^H^ increases to almost one (Figure 3b). This data indicates that Asp226 accepts a proton first as pH decreases, which is in agreement with its macroscopic p*K*_a_ (5.3) being higher than Asp38 (3.2).

To corroborate the results, we performed another set of replica-exchange CpHMD simulations starting from a different crystal structure (PDB ID: 3sfc,^31^ inhibitor removed). These simulations gave similar results. The residue-specific p*K*_a_’s are 3.5 and 5.8 for Asp38 and Asp226, respectively, and the stepwise p*K*_a_’s are 3.3 and 5.9 attributable to Asp38 and Asp226, respectively (Table 1, Figure S2).

**Table 1.**
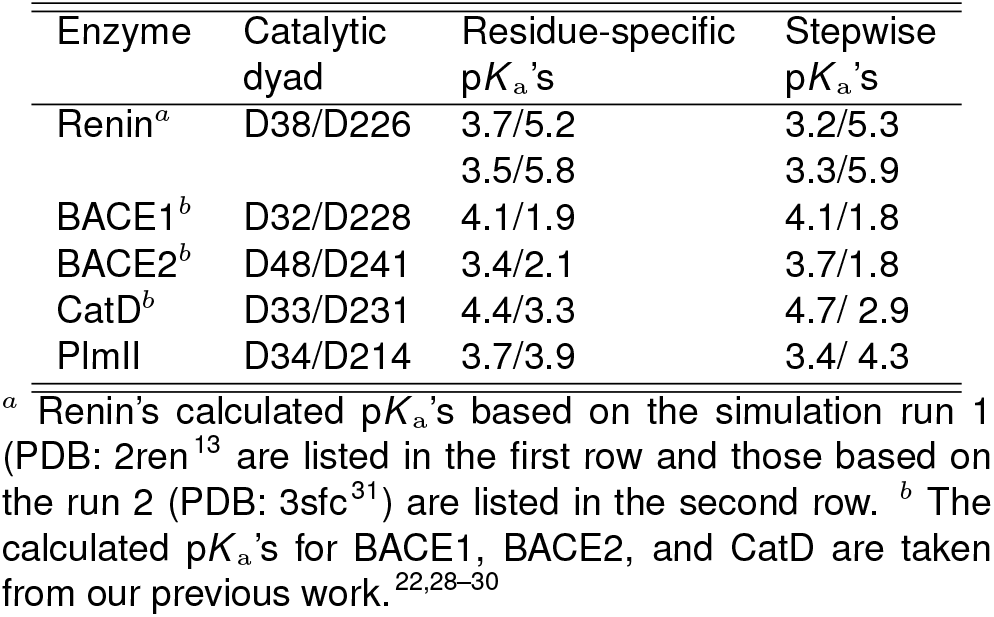
Calculated p*K*_a_âĂŹs of renin’s catalytic dyad in comparison to those of homologous pepsin-like aspartic proteases.

| Enzyme | Catalytic dyad | Residue-specific $pK_a$'s | Stepwise $pK_a$'s |
| --- | --- | --- | --- |
| Renin<sup>a</sup> | D38/D226 | 3.7/5.2<br>3.5/5.8 | 3.2/5.3<br>3.3/5.9 |
| BACE1<sup>b</sup> | D32/D228 | 4.1/1.9 | 4.1/1.8 |
| BACE2<sup>b</sup> | D48/D241 | 3.4/2.1 | 3.7/1.8 |
| CatD<sup>b</sup> | D33/D231 | 4.4/3.3 | 4.7/ 2.9 |
| PlmII | D34/D214 | 3.7/3.9 | 3.4/ 4.3 |

<sup>a</sup> Renin's calculated $pK_a$ 's based on the simulation run 1 (PDB: 2ren<sup>13</sup>) are listed in the first row and those based on the run 2 (PDB: 3sfc<sup>31</sup>) are listed in the second row. <sup>b</sup> The calculated $pK_a$ 's for BACE1, BACE2, and CatD are taken from our previous work.<sup>22,28–30</sup>

### Number of hydrogen bonds is the major determinant of the dyad p*K*_a_ order

Solvent accessibility, hydrogen bonding, and Coulomb interactions are three major factors contributing to the p*K*_a_ shift of an amino acid sidechain relative to the solution or model value. Catalytic residues are typically buried in the protein interior which lacks of other charged/titratable residues. Our previous work based on CpHMD simulations of several enzymes, BACE1/BACE2, capthepsin D, hen egg lysozyme, and Staphylococcal nuclease showed that solvent accessibility and hydrogen bonding are the determinants of the p*K*_a_ order for the catalytic dyad. ^22,29,30^ We found that both dyad residues are acceptors to a number of hydrogen bonds formed during the MD simulations, and the general base (or nucleophile) which has a lower p*K*_a_ forms a significantly larger number of hydrogen bonds than the general acid (or proton donor). ^22,29,30^ We also found that the nucleophile is more solvent exposed as compared to the proton donor. ^22,29,30^ These two observations can be rationalized, as the formation of hydrogen bonds stabilizes the deprotonated aspartate, while solvent exclusion stabilizes the protonated form.

To test the hypothesis that the general base Asp38 is more solvent accessible than Asp226, we calculated the number of water (hydration number) surrounding each dyad residue as a function of pH. As expected, the hydration number for both Asp38 and Asp226 increases with pH, indicating that water enters the active site as the degree of deprotonation increases (Figure 3c). Closer examination revealed that the hydration increase starts at around pH 4 and plateaus at pH 6–7, which coincides with the decrease in the fraction of the singly protonated state D38^−^/D226^H^ and with the increase in the fraction of the doubly deprotonated state. In roughly the same pH range, the number of bridging water between the two residues increases from one and then plateaus at about 1.5 (Figure 3c). In the entire pH range, the hydration number for Asp226 is slightly larger than Asp38, although the difference is very small. Thus, the data of renin does not support the hypothesis that the general base is more hydrated.

Next, we tested the hypothesis that the deprotonated Asp38 forms a larger number of hydrogen bonds than the proton donor Asp226. Below pH 7, Asp38 forms up to one more hydrogen bond than Asp226, while above pH 7 when both residues are fully deprotonated, the number of hydrogen bonds is comparable (Figure 3d). Importantly, the pH profile of the hydrogen bond count for each residue matches the corresponding titration curve. As Asp38 deprotonates in the pH range 2–5, its hydrogen bond count increases from 1 to 1.5; as Asp226 deprotonates in the pH range 4-8, its hydrogen bond count increases from 0.5 to 1.5. This data supports the hypothesis that the Asp38 forms more hydrogen bonds than Asp226, suggesting that it is the driving force for the lower p*K*_a_ of Asp38 relative to Asp226.

### Detailed hydrogen bond environment of the aspartyl dyad

To further understand the physical origin of the different proton affinities of the dyad residues, we examined the specific hydrogen bonds (Figure 4). The carboxylate groups of Asp38 and Asp226 are within hydrogen bonding distance of each other in the crystal structure (PDB: 2ren^13^). Simulations showed that the D38–D226 hydrogen bonds are formed in a pH-dependent manner in the pH conditions where at least one of the two residues samples the protonated state (pH < 6), indicating that it serves as the proton donor while the other serves as the proton acceptor.

**Figure 4.**
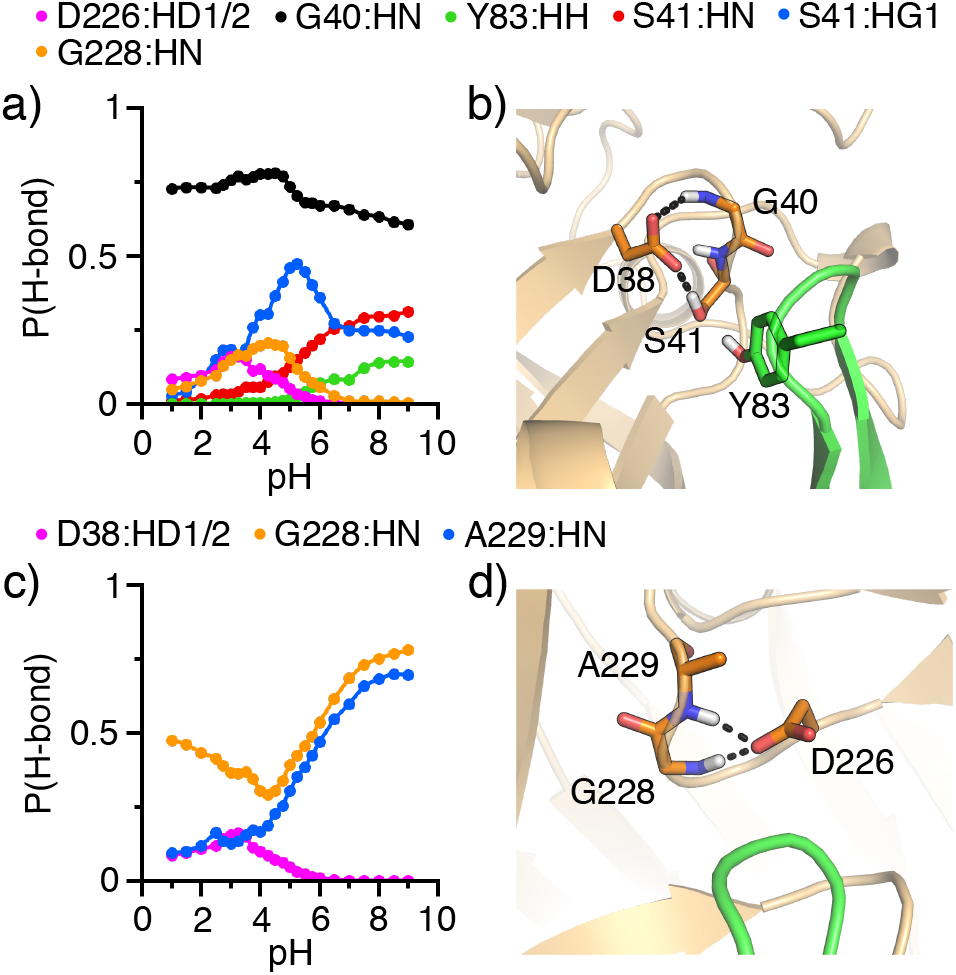
Hydrogen bond formation of the catalytic dyad in renin. **a), c)** Probabilities of forming specific hydrogen bonds with the carboxylates of Asp38 (**a**) and Asp226 (**c**) as a function of pH. Proton donors are indicated in different colors and shown in the legend. **b), d)** Snapshots showing the hydrogen bond environment of Asp38 (**b**) and Asp226 (**d**). Data from the simulation run 1 was used.

In addition to the interaction with Asp226, the carboxylate of Asp38 accepts hydrogen bonds from several other residues, including Gly40, Ser41, Tyr83, and Gly228 (Figure 4a and b). Asp38 maintains a stable hydrogen bond with the backbone amide of Gly40 in the entire simulation pH range, and forms hydrogen bonds with the backbone amide and sidechain hydroxyl of Ser41. Interestingly, while the hydrogen bond with the backbone Ser41 becomes more stable with pH (probability increases), the hydrogen bond with the sidechain Ser41 is most stable between pH 4 and 7 and the probability reaches a maximum around pH 5.5.

In addition to the interaction with Asp38, the carboxylate of Asp226 is a hydrogen bond acceptor to Gly228 and Ala229 (Figure 4c and d). Interestingly, Gly228 can form hydrogen bonds with both dyad residues. While the pH profile for the Gly228–Asp38 hydrogen bond is bell shaped with a maximum round pH 4, the pH profile for the Gly228–Asp226 hydrogen bond has an inverted bell shape with a minimum at the pH 4. This behavior can be readily understood from the difference in the titration pH range for the two residues. In the pH range 2–4, the probability for the Gly228–Asp38 hydrogen bond increases with pH due to the increasing deprotonation of Asp38 while Asp226 remains fully protonated. Above pH 4, the probability for the Gly228–Asp226 hydrogen bond increases due to the increasing deprotonation of Asp226 while Asp38 remains fully deprotonated. Above pH 4, Asp226 also accepts a hydrogen bond from the backbone amide of Ala229, and the probability increase with pH.

### Comparison to the pH-activity measurement

The order of the calculated macroscopic p*K*_a_’s of 3.2(3.3) and 5.3(5.9) attributable to Asp38 and Asp226, respectively, is in agreement with the pH-dependent activity measurements of human renin with two substrate peptides.^19^ The macroscopic p*K*_a_’s of 5.3 and 6.3 were obtained in the presence of the peptide PTDP, which represents the wild-type porcine angiotensinogen (P1–P4 sequences are the same as the human one), while the p*K*_a_’s of 4.4 and 7.4 obtained for the mutant peptide HP2Q, in which the P2 His is substituted with Gln.^19^ The authors suggested^19^ that both Asp38 and Asp226 in the apo renin are deprotonated at physiological pH, and during catalysis unprotonated Asp38 is the base that abstracts a proton from the nearby catalytic (nucleophilic) water, while Asp226 forms a charged hydrogen bond (salt bridge) with the P2 His, allowing the pair to form the “surrogate” acid (proton donor) at physiological pH.

The simulation data of the apo renin are consistent with the above experiment. In the simulations, a bridging water between the dyad residues is always present (Figure 3c), which is consistent with presence a catalytic water as proposed by experiment.^19^ The lower p*K*_a_ of Asp38 relative to Asp226 is in agreement with the respective roles of general base and acid as suggested by the experiment.^19^ Our calculated dyad p*K*_a_’s of 3.2(3.3)/5.3(5.9) are lower than the experimental estimate of 4.4/7.4 (without interaction with the P2 His) for the apo renin. This discrepancy is consistent with our previous CpHMD titration studies of aspartyl proteases which showed a systematic underestimation of 1–1.5 pH units,^22,28^ likely due to the limitation of the hybrid-solvent method. The lower p*K*_a_’s compared to experiment may also be attributed to the fact that the active site in the simulations is more hydrated than the substrate-bound state observed in experiment.

### Conformational dynamics of the flap

A common structural feature of aspartyl proteases is a *β*-hairpin loop that covers the active site, commonly known as the flap.^33,34^ Our previous CpHMD simulations showed that the flap in BACE1 is flexible and can open and close in a pH-dependent manner.^28^ At low and high pH, the flap in BACE1 is closed, while at intermediate pH where the enzyme is active (pH 3.5–5.5), the flap is open.^28^ In contrast, CpHMD simulations showed that the flap in CatD is rigid and remains open in a broad pH range 2.5–6, consistent with the pH activity profile.^29^ Thus, we were curious about the conformational dynamics of renin’s flap and whether it is pH dependent. To examine the position of renin’s flap relative to the dyad, we used two distances, R1, the distance between Tyr83:OH and Asp38:CG and R2, the distance between Ser84:CB and Asp226:CG (Figure 5a). Tyr83 is a conserved residue among all pepsin-like aspartyl proteases and is capable of forming a hydrogen bond with the dyad in BACE1 to prevent substrate access.^28,35^ Ser84 is located near the tip of the flap and other aspartyl proteases has a similar amino acid at the same location (Figure 2).

**Figure 5.**
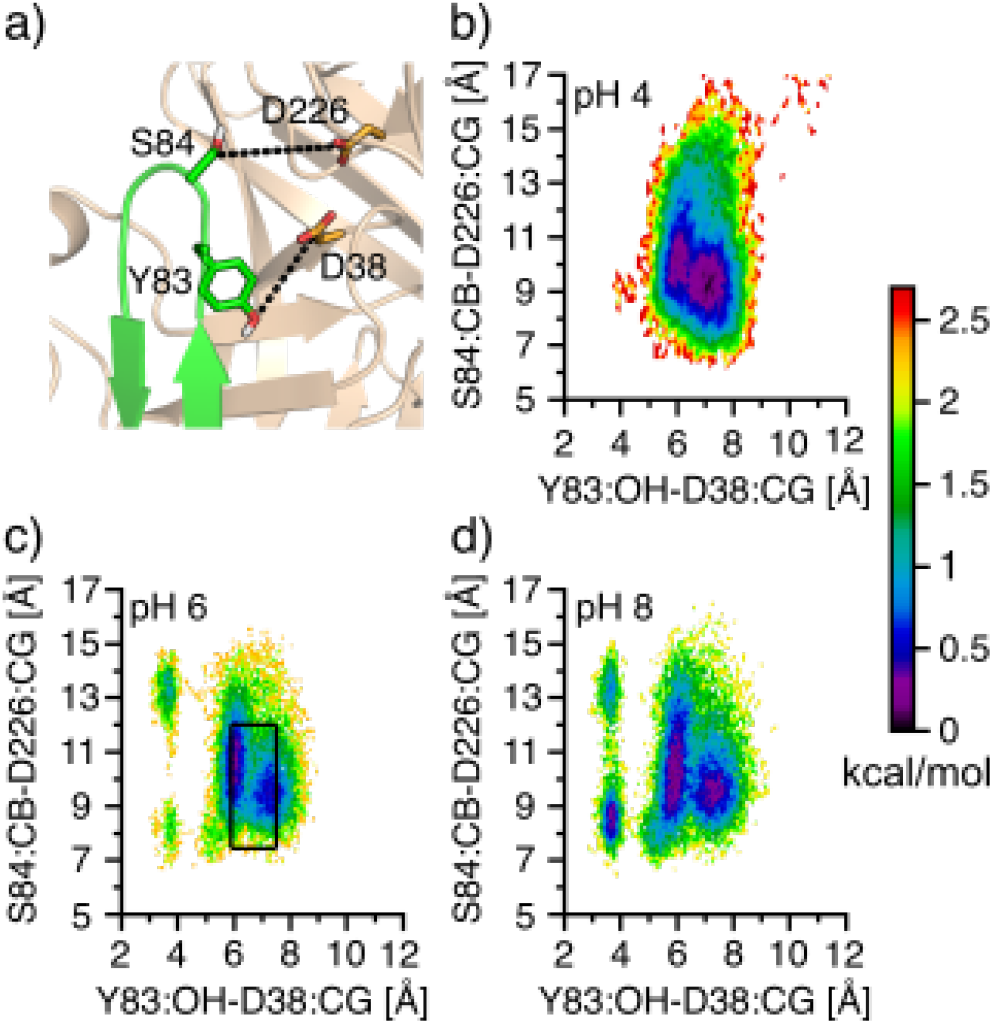
Conformational dynamics of the flap in renin. a) Zoomed-in view of the flap region. The two distances used to calculate the free energy surface are indicated as dashed lines. b–d) Free energy surface as function of the distance between Tyr83:OH and Asp38:CG and the distance between Ser84:CB and Asp226:CG from the simulations at pH 4 (b), 6 (c), and 8 (d). The black box indicates the distances sampled by the majority of the crystal structures (see Figure S5). Data from the simulation run 1 was used.

The free energy surface as a function of R1 and R2 was calculated for different pH conditions (Figure 5b-d and Figure S3). At all pH conditions, renin’s flap samples a broad free energy minimum region with R1 between 5 and 7.5 Å and R2 between 7 and 12 Å; however, at pH greater than 5.5, two additional minima appear with R1 located at 3.5 Å and R2 located at 8 or 13 Å. Close examination revealed that these two minima represent the conformations in which the hydroxyl group of Tyr83 donates a hydrogen bond to the carboxylate of Asp38. An analogous hydrogen bond was found for BACE1 ^28,35^ and other aspartyl proteases,^36,37^ and its formation is linked to the flap closure, leading to the so-called Tyr-inhibited state. Our analysis shows that the occupancy of the Tyr83–Asp38 hydrogen bond in renin increases between pH 5 and 7.5 and plateaus at about 20% (Figure S4), which is significantly lower than the analogous Tyr-Asp hydrogen bond in BACE1.^28^ It is worth noting that, despite the Tyr-Asp hydrogen bond, the tip of renin’s flap can sample two positions, with the R2 value of 8 and 13 Å, respectively, suggesting that the flap remains flexible and open. This is in stark contrast to the flap of BACE1, as the analogous Tyr is near the tip of the flap, and its hydrogen bonding with the catalytic Asp forces the flap into a closed state.

### Comparison of the flap conformation with that observed in the X-ray crystal structures

We compare renin’s flap dynamics observed in the simulations with the available crystal structures of renin in complex with substrate or inhibitor. A scatter plot of 86 PDB entries (165 subunits) of human renin structures (pH conditions vary from 3.0 to 8.5) shows that in a majority of structures, the Tyr83–Asp38 distance falls into the range of 5.8–7.5 Å and the Ser84-Asp226 distance falls into the range of 7.4–12 Å. Thus, the free energy minimum region from the simulations is in agreement with the majority of the X-ray structures (Figure 5c, black box). Curiously, a few crystal structures show the Tyr83–Asp38 distance above 8.9 and the Ser84-Asp226 distance above 13 Å (Figure S5). These structures correspond to those with large inhibitors (with 38 to 44 heavy atoms), which break the close contact between Trp45 and Tyr83, forcing the flap to a widely open conformation. Interestingly, this flap conformation is occasionally sampled at pH 4 (Figure 5b) but missing at higher pH (Figure 5c-d). It is worth noting that the Tyr-inhibited state observed in the simulations is not represented by any crystal structures (Figure 5c and Figure S4), which is perhaps due to the fact that most crystal structures except for two are inhibitor/substrate-bound complexes.

### Comparison of the dyad protonation states with other pepsin-like aspartyl proteases

We compare the calculated dyad p*K*_a_’s of renin with those of the homologous pepsin-like aspartic proteases BACE1, ^28^ BACE2, ^22,30^ and CatD,^29^ which share a sequence similarity level of 43%, 47%, and 66%, respectively, with renin (Figure 1). Surprisingly, the p*K*_a_ order of renin’s catalytic dyad is opposite to that of BACE1, BACE2, and CatD. According to the CpHMD simulations and experiment,^22,28–30^ the residue homologous to Asp38 in renin, Asp32 in BACE1, Asp48 in BACE2, or Asp33 in CatD, has a higher p*K*_a_ than the residue homologous to Asp226 in renin, Asp228 in BACE1, Asp241 in BACE2, or Asp231 in CatD (Table 1). The relative p*K*_a_’s suggest that Asp32, Asp48, and Asp33 are protonated serving the role of a general acid, while Asp228, Asp241, and Asp331 are deprotonated serving the role of a general base in the catalytic reactions of BACE1, BACE2, and CatD, respectively.

Interestingly, the relative p*K*_a_’s of renin is identical to those of the pepsin-like protease PlmII which shares 53% sequence similarity with renin (Table 1). The CpHMD titration showed that Asp34 and Asp214 in PlmII, which are analogous to Asp38 and Asp226 in renin, have the stepwise p*K*_a_’s of 3.4 and 4.3, and thus serving the roles of general base and general acid, respectively. We note, a detailed study of PlmII will be published in a future work. Taken together, the comparison shows that the catalytic roles of the dyad are not conserved among the pepsin-like proteases.

## CONCLUSION

CpHMD simulations have been performed on the apo renin to understand the roles of the catalytic dyad and conformational dynamics of the flap. The calculated macroscopic p*K*_a_’s attributable to Asp38 and Asp226 from two sets of CpHMD simulations are 3.2(3.3) and 5.3(5.9), respectively, suggesting that Asp38 serves as a general base and Asp226 serves as a general acid during renin catalysis. Interestingly, the relative p*K*_a_ order of the catalytic dyad in renin are opposite to BACE1, BACE2, CatD, but identical to PlmII, suggesting that the acid/base roles are not conserved among pepsin-like aspartyl proteases. The analysis of CpHMD trajectories shows that the deprotonated carboxylate of Asp38 forms more hydrogen bonds than Asp226, consistent with our previous finding that the general base (or nucleophile) of a catalytic dyad accepts more hydrogen bonds than the general acid (or proton donor) and that some of the hydrogen bonds are absent in the X-ray crystal structure but emerge during the proton-coupled conformational sampling in the simulation. ^22^

The catalytic roles assigned by the CpHMD simulations are in agreement with those based on the pH-activity measurement.^19^ The latter yielded Asp38/Asp226 p*K*_a_’s of 5.3/6.3 in the presence of the wild-type substrate and 4.4/7.4 in the presence of a mutant substrate, in which the P2 His was substituted by Gln. The differences in the p*K*_a_’s were hypothesized to originate from a charged hydrogen bond between Asp226 and the P2 His, which allows the Asp226–His ion-pair to act as a general acid in catalysis at neutral pH.^19^ The Asp226–His interaction was also hypothesized to raise the p*K*_a_ of Asp38, thus shifting the enzyme optimum pH higher to the neutral range.^19^ Considering the known systematic underestimation of 1–1.5 pH units for the aspartyl protease dyad p*K*_a_’s by hybrid-solvent CpHMD,^22,28^ our calculated p*K*_a_’s of 3.2(3.3)/5.3(5.9) for Asp32/Asp226 are consistent with the experimental estimate of 4.4/7.4 for the apo renin, thus supporting the hypothesis that renin’s higher optimum pH relative to other pepsin-like proteases is due to the effect of substrate interactions. To directly test the hypothesis of substrate-directed catalysis, future simulations will be carried out using the substrate-bound renin structure.

The CpHMD simulations also revealed the distinctive flap dynamics in renin. Unlike the flap in BACE1 which displays pH-dependent dynamics, renin’s flap is open regardless of pH, similar to BACE2 and CatD. However, the Tyr-inhibited state, in which the conserved Tyr83 forms a hydrogen bond with Asp38, is occasionally sampled above pH 5 and the population increases to 20% at pH 7.5; this state is not sampled by BACE2 and CatD. Interestingly, while the analogous hydrogen bond in BACE1 prevents the flap from opening, the flap in renin remains open, as Tyr83 is positioned lower on the flap and not on the tip as in BACE1. Aspartyl proteases are an important class of enzymes; our work demonstrates that CpHMD simulations is a powerful tool for advancing the detailed knowledge of their pH-dependent structure-function relationships which remain poorly understood.

## METHODS and PROTOCOLS

### System preparation

The coordinates of the X-ray crystal structures of human renin (PDB: 2ren,^13^ apo) and an inhibitor-bound complex (PDB: 3sfc,^31^ subunit B) were retrieved from the PDB. The positions of the missing residues 53-55, 166-170, and 287-295 in the apo renin structure (2ren) were built by superimposing the backbone atoms onto those of the holo structure (3sfc) which has coordinates for all residues. The root-mean-square deviation (RMSD) of the backbone atoms between the two structures was 0.84 Å. For PlmII, the X-ray crystal structure (PDB: 1sme^32^) was used with the ligand being removed. The hydrogen atoms were added using the HBUILD facility in CHARMM.^38^ To remove unfavorable contacts, the apo renin structure was first minimized 100 steps using the Adopted Basis Newton-Raphson (ABNR) method, and the PlmII structure was energy minimized using 10 steps of steepest descent (SD) and 10 steps of Adopted Basis Newton-Raphson (ABNR) method. The protein was then solvated in an octahedral water box with a heavy-atom distance of at least 10 Å between the protein and the edge of the water box. Following solvation, the water positions were energy minimized in several stages. First, 50 steps of SD followed by 50 steps of ABNR minimization was performed, with the protein heavy atoms fixed. Next, a five-stage restrained minimization was performed, where the harmonic force constant on the backbone heavy atoms was 100, 50, 25, 5, and 0 kcal·mol^−1^·Å^−2^. Each stage included 50 steps of SD and 100 steps of ABNR, except for the first stage which included 50 steps of SD and 10 steps of ABNR minimization.

Two simulations of renin were performed; run 1 started from the apo structure and run 2 started from the holo structure with the inhibitor removed. Only one simulation was performed for PlmII. All simulations were performed with the CHARMM program c36a2. ^38^ The protein was represented by the CHARMM22/CMAP all-atom force field,^39^,^40^ and water was represented by the modified TIP3P water model.^38^ The system was gradually heated from 100 K to 300 K over the course of 120 ps. The system was subsequently equilibrated for 280 ps in four stages, where the harmonic force constant for the protein heavy atoms was 5 (40 ps), 1 (40 ps), 0.1 (100 ps), and 0 (100 ps) kcal·mol^−1^ ·Å^−2^ (100 ps). The system was further equilibrated for 580 ps without any restraint. In the heating and equilibration stages, constant pH functionality (PHMD module in CHARMM) was turned on and pH was set to the crystal pH conditions (pH 4.7 for the renin simulation run 1, pH 4.5 for the renin simulation run 2, and pH 6.5 forthe plmII simulation).

### Replica-exchange CpHMD simulations

Following equilibration, hybrid-solvent CpHMD simulations with the pH replica-exchange protocol were performed. Detailed methodology can be found in the original work^26^ and a review.^27^ 24 replicas were placed in the pH range 1-9 (1-8.5 for plm II). Each replica was simulated in the NPT ensemble at 300 K and 1 atm. The particle mesh Ewald method^41^ was used to calculate long-range electrostatic interactions, with a real space cutoff of 12 Å and a sixth-order interpolation with a 1-Å grid spacing. The SHAKE algorithm was used to constrain bonds involving hydrogen atoms to enable a 2-fs timestep. A Generalized Born (GB) calculation was invoked every 10 molecular dynamics (MD) steps to update the titration coordinates. Every 500 MD steps (1 ps), the adjacent pH replicas attempted to swap conformational states based on the Metropolis criterion. ^26^ All sidechains of Asp, Glu, and His residues were allowed to titrate. Each replica ran for 28 and 27 ns in the renin simulation run 1 and 2, with the aggregate sampling time of 672 ns and 648 ns, respectively, In the PlmII simulation, each replica ran for 29 ns, with the aggregate time of 696 ns. To verify convergence, the time series of the cumulatively calculated p*K*_a_ values of the catalytic dyad were examined (Figures S1). For analysis, the data from the first 5 ns per replica was discarded for renin and the first 9 ns was discarded for PlmII.

### p*K*_a_ calculations

In CpHMD simulations,^27^ the continuous variables λ and *x* are used to represent the titration coordinates and tautomer interconversion, respectively. A protonated state was defined as those with λ < 0.2, and *x* < 0.2 or x > 0.8, while an unprotonated state was defined as those with λ > 0.8, and x < 0.2 or x > 0.8. Accordingly, the fraction of unprotonated state (*S*) was calculated for each titratable site at each simulation pH. The microscopic residuespecific p*K*_a_ was calculated by fitting *S* at different pH to the generalized Henderson-Hasselbalch (modified Hill) equation,

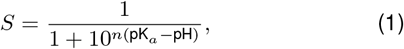

where *n* is the Hill coefficient that represents the slope of the transition region in the titration curve. To compare with experiment, we also calculated the macroscopic stepwise p*K*_a_’s by fitting the total number of bound protons to the catalytic dyad (*N*_prot_) to the following statistical mechanics based two-proton model:^42^,^43^

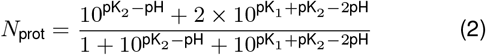

where pK_1_ and pK_2_ are the macroscopic p*K*_a_’s of the dyad, and the denominator represents the partition function.

## Supporting Information Available

Supporting Information contains supplemental tables and figures.

## Acknowledgement

Financial support from the National Institutes of Health (GM09888) is acknowledged. S. Ma would like to thank Dr. Cheng-Chieh (Kevin) Tsai, Dr. Zhi Yue, and Dr. Robert Harris for their help in setting up CpHMD simulations and data analysis.

