## Supplemental figures for "Exploring the pH-dependent structure-dynamics-function relationship of human renin"

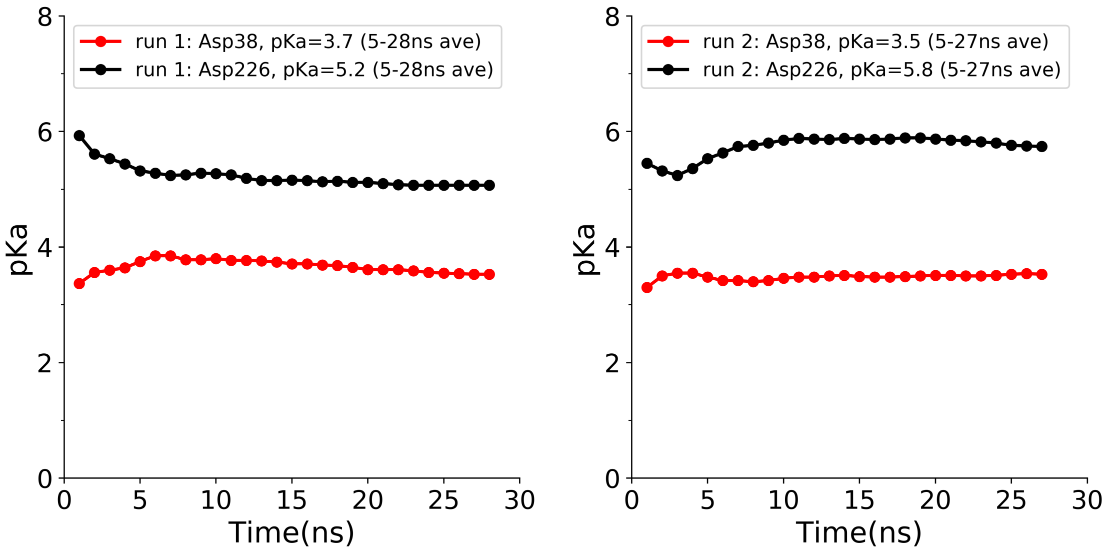


Figure S1: **Convergence of the calculated residue-specific p$K_a$'s of the catalytic dyad in human renin.** Left. Simulations starting from the X-ray structure PDB 2ren (run 1). Right. Simulations starting from the X-ray structure PDB 3sfc (run 2). The data for Asp38 and Asp226 are shown in red and black, respectively.

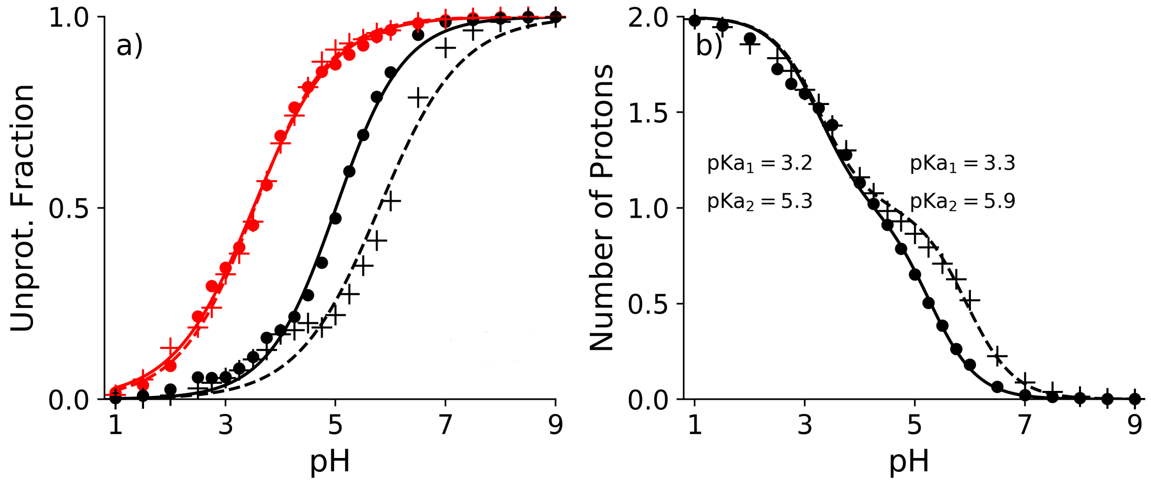


Figure S2: **Microscopic and macroscopic titration curves of the catalytic dyad in human renin.** a) The unprotonated fractions of Asp38 (red) and Asp226 (black) as a function of pH calculated from the simulations with the structures 2ren (run 1, dots) and 3sfc (run 2, crosses). The best fits to the generalized Henderson-Hasselbalch equation are given (solid lines for 2ren and dashes lines for 3sfc). b) The number of protons for the catalytic dyad calculated from the simulations with the structures 2ren (run 1, dots) and 3sfc (run 2, crosses). Data from the first 5 ns per replica in both simulations were discarded in the calculations.

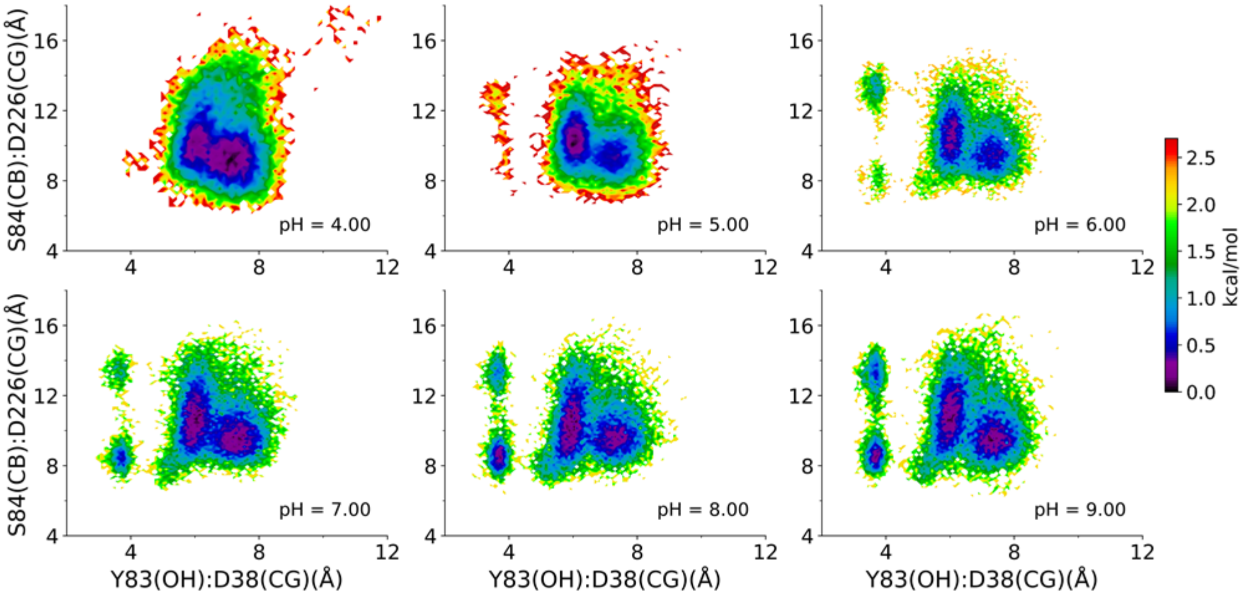


Figure S3: Free energy surfaces as a function of the distance between Tyr83:OH and Asp38:CG and the distance between Ser84:CB and Asp226:CG at pH 4–9. Data from the simulation run 1 were used.

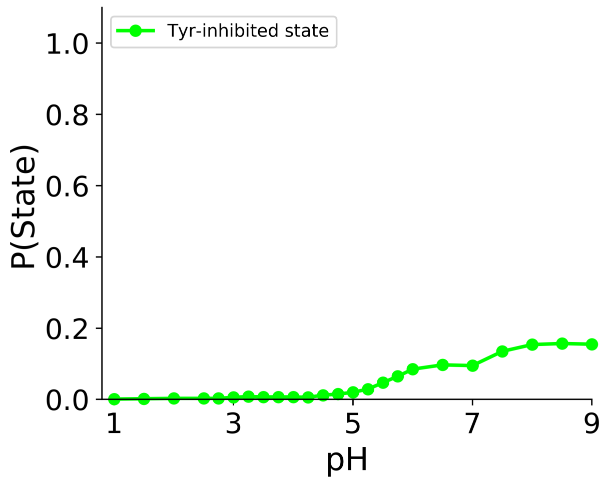


Figure S4: Probability of the Tyr-inhibited state: R1 < 5.0 Å; Where R1: Tyr83(OH)-Asp38(CG).Data from the simulation run 1 were used.

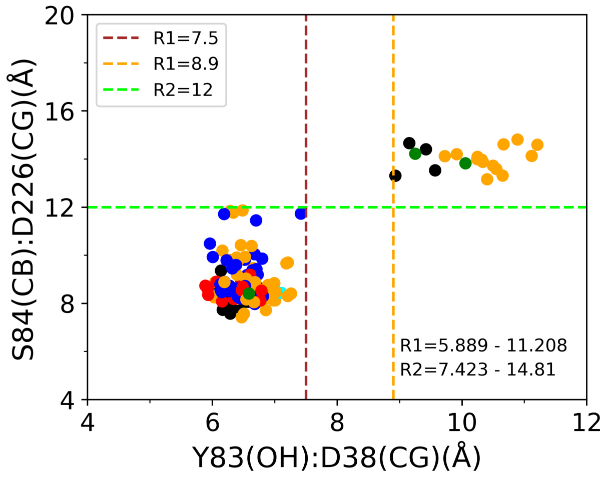


Figure S5: The distance between Tyr83:OH and Asp38:CG (R1) vs. the distance between Ser84:CB and Asp226:CG (R2) in the crystal structures of human renin (apo or bound to substrate mimetic peptides or small molecule inhibitors). Data are color coded according to the crystallization pH condition: red (pH 3-4), orange (pH 4-5), cyan (pH 5-6), green (pH 6-7), blue (pH 7-8.5), and black (pH not provided in the PDB file).
